# Structural and functional role of residues undergoing hereditary spastic paraplegias-linked mutations: insights from a simulation of the spiral to ring transition in katanin

**DOI:** 10.1101/2024.11.27.625675

**Authors:** Maria S. Kelly, Riccardo Capelli, Ruxandra I. Dima, Paolo Carloni

## Abstract

Several dozen mutations in the large human spastin enzyme assembly have been associated with mobility impairment in hereditary spastic paraplegias. Some of them impact the structural determinants of two functional conformations of the protein, spiral and ring. Here we investigate the possible effect of the mutations on the transition between the two conformations, which is essential for the enzymatic function. By performing a variety of molecular simulations (including metadynamics) on the closely related protein katanin, we suggest that about one fourth of the known disease-associated mutations affect the transition and/or the stability of a previously unrecognized intermediate. The protocol used here can be applied to the study of conformational changes in other large biomolecular complexes.

**Significance Statement:** By combining molecular dynamics and enhanced sampling simulations with custom-designed collective variables, we explore transient intermediate states in a large protein assembly, the severing enzyme katanin. This enzyme plays a critical role in neuronal microtubule remodeling. By targeting the transition between the spiral and ring conformations of the protein, we reveal mechanisms associated with neurodegenerative mutations that impair the function of spastin, a structurally and functionally similar enzyme. These findings provide insights that may inform future therapeutic strategies. They also expand our ability to simulate molecular processes relevant to health and disease, as our approach can be readily applied to study structural changes in other complex biological systems.

---

**H**uman spastin is a microtubule severing enzyme that is essential for shaping the microtubule network within neurons, particularly in axon growth and maintenance (1, 2). The protein removes tubulin dimers from microtubules in order to modify their lattice (3). Dysfunction of the protein is associated with neurodegenerative disorders that can lead to partial or complete loss of mobility of humans, named Hereditary Spastic Paraplegias (HSPs) (4–8). As many as 65 missense mutations have been associated with the disease (Tab. S1) (9). Some of them affect spastin’s ability to bind ligands and oligomerize into its functional quaternary structure (Tab. S1) (9).

Human spastin belongs to the ATPases Associated with diverse cellular Activities (AAA+) superfamily of proteins. It shares function and most of its structural determinants with the AAA+ protein katanin (10–13): **(i)** The *nucleotide-binding domain* (NBD), responsible for ATP binding and hydrolysis (ATPase motor). As shown by cryoelectron microscopy (cryo-EM, Fig. 1), the ATP-binding pocket is formed by the *Walker A, Walker B*, and *Arg-finger* motifs. The cofactor is stabilized by a Mg(II) ion. The NBD interacts with the C-terminal tails of tubulin (CTTs). These are directly exposed to the mechanical forces generated by the enzymes, leading to the severing of the microtubules (2, 11). Two positively charged pore loops (PL1 and PL2), located in the NBD and protruding into the central pore, form a spiral around the negatively charged CTT of the microtubules (required for substrate processing, Fig. 1). Residues in the pore loop 3 (PL3) form hydrogen bonds with both the nucleotide and PL2, thus connecting ATP and CTT binding (Fig. 1).^∗^ **(ii)** The *microtubule-interacting and trafficking* (MIT) domain, formed by a three-helix bundle. This provides anchoring support for the motor to make contact with the microtubules, and a flexible linker region that connects the MIT with the NDB (10, 14), enabling proper orientation of the latter on the microtubule lattice (Fig. 1). **(iii)** The *Helical Bundle Domain* (HBD), unique for these two proteins across the AAA+ family (Fig. 1). The HBD of each protomer binds to the NBD of the neighboring protomer, forming an asymmetric spiral hexamer with a large gate between the two terminal protomers (Fig. 2) (11, 12, 15). This opening allows entrance of the CTT into the hexamer’s pore where the PLs are located (Fig. 2).

**Fig. 1.**
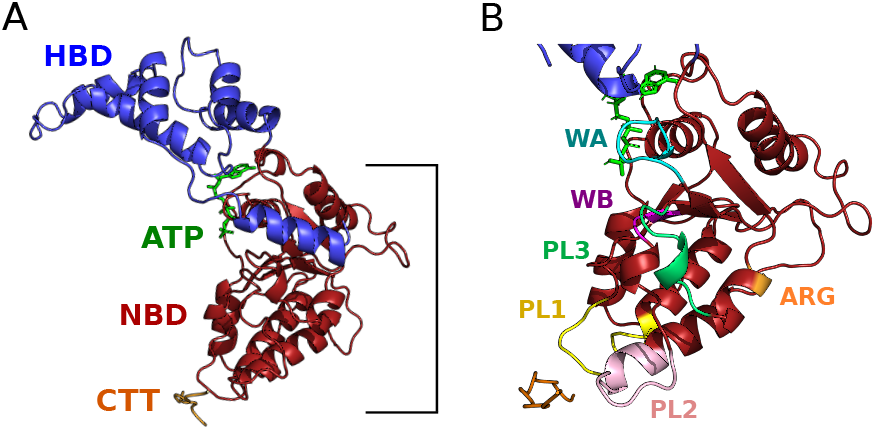
(A) Katanin monomer with domains (NBD and HBD) and ligands (ATP and CTT) labeled in their respective colors. The details of the binding site, with the Mg(II) are shown in Fig. S1. (B) Katanin’s NBD domain with functional motifs colored and labeled. C-terminal helix in HBD was removed for visual aid.

**Fig. 2.**
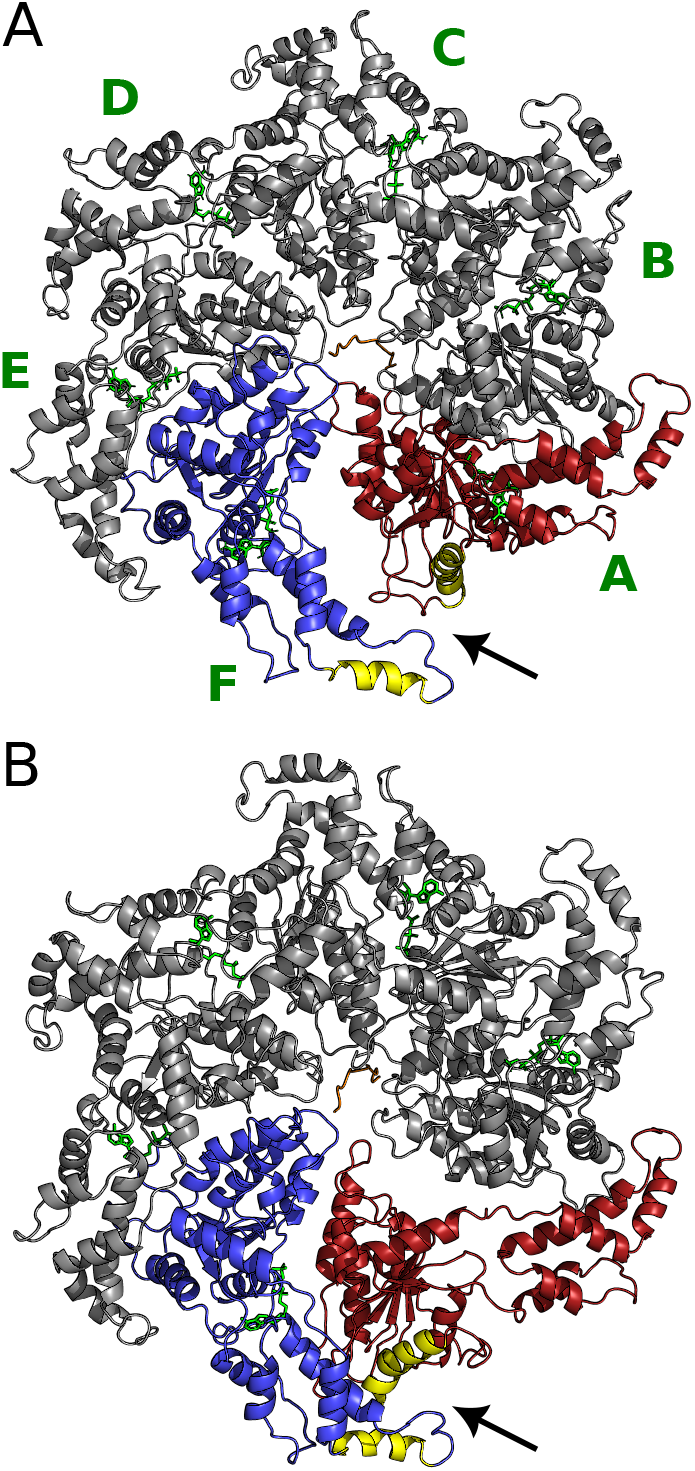
Katanin spiral (A) and ring (B) conformers. The protomers A-F are labeled. The terminal protomers A and F colored in red and blue, respectively. The two alpha helices used for CV_1_ are colored in yellow. Protomers A-F are labeled in the spiral. The HBD - NBD interaction and the entrance of the CTT into the hexamer’s pore are shown with an arrow. The MIT domains were unresolved in the cryo-EM structures and are therefore not depicted here.

The ATP hydrolysis in one of the two terminal NDBs (Fig. 2) leads to a transition of the spiral structure to its ring structure (S → R, Fig. 2), where loss of the nucleotide within protomer A results in the closure of the terminal gate.

This is an essential part of the severing mechanism as it allows the generation of a force on the microtubule lattice needed to extract the tubulin dimers (1). An important question is the following: do any of the disease-linked mutations affect this spiral-ring transition? To address this issue, here we uncover the structural determinants of the S → R transition - so far unknown - using molecular simulations. We focus on katanin from *C. elegans* because of the availability of much larger structural information on this enzyme than for human spastin, the structural similarities with spastin, their similar functional role, and the large of number of conserved residues undergoing HSPs-linked mutations (See Supporting Information for details).

Given the large size of the hexameric complex (consiting of 1,910 aminoacids), we first investigate qualitatively the S → R transition in katanin using an efficient non-equilibrium technique, ratchet&pawl MD(18, 19)- similar to steered molecular dynamics (20)- as a function of two apt collective variables (CVs): (i) the distance between two helices of opposite terminal protomers to control the opening and closing of the S → R transition, and (ii) a linear combination of distances between pairs of atoms forming intermolecular interactions (salt bridges and hydrogen bonds) in the ring and spiral states, computed by the dimensionality reduction technique *Harmonic Linear Discriminant Analysis* (HLDA)(21, 22). Next, we use Well-Tempered Metadynamics (WT-MetaD), an accurate technique able to explore the free energy landscape, as a function of the two CVs (23, 24). We exploit the ability of this technique to escape free energy minima to find out-of-pathway relevant transients (25). Indeed, the simulations successfully identify an intermediate state that rationalizes the mechanistic effects of several disease-related residues, identifying them as essential for the severing event to take place. This protocol is applicable to large real-world systems, where exhaustive exploration by normal enhanced sampling approaches is out of reach even with current computational capabilities.

## 1. Results

We identified two CVs able to describe the S → R transition in katanin by monitoring selected distances in 150 ns MD simulations of both spiral and ring conformers. CV_1_ was defined as the distance between the centers of mass of specific helices from protomer A and protomer F (Fig. 2). The helices were selected because they were the most affected by the S → R transition, assuring that CV_1_ results informative in tracking the passage from one state to the other. CV_2_ incorporates essential changes of the intramolecular interactions involving the three PLs. As discussed in the Introduction, these are functionally relevant to substrate processing (10, 11). To distill such information, we aggregate the descriptors linked to the nonbonded interactions (*i.e*., the distances between pairs of atoms involved in salt bridges and hydrogen bonds) as in ref. (25): CV_2_ is then the 1st eigenvector generated from the HLDA (a linear combination of distances. see Tab. S2) that best distinguishes the spiral conformer from the ring one. Almost all (97%) of the identified pairs match with corresponding spastin residues labeled as disease-related, further confirming the similarity between the two proteins (3).

Next, we tested the ability of the two CVs to describe the S → R transition by performing CV-based Ratchet&Pawl MD simulations (18, 19). These simulations led satisfactorily to both forward and backward S → R transitions in a very short time (less than 100 ns), suggesting that the two chosen CV are appropriate to study this process. The conformational space sampled in this simulation is shown in Fig. 3.

**Fig. 3.**
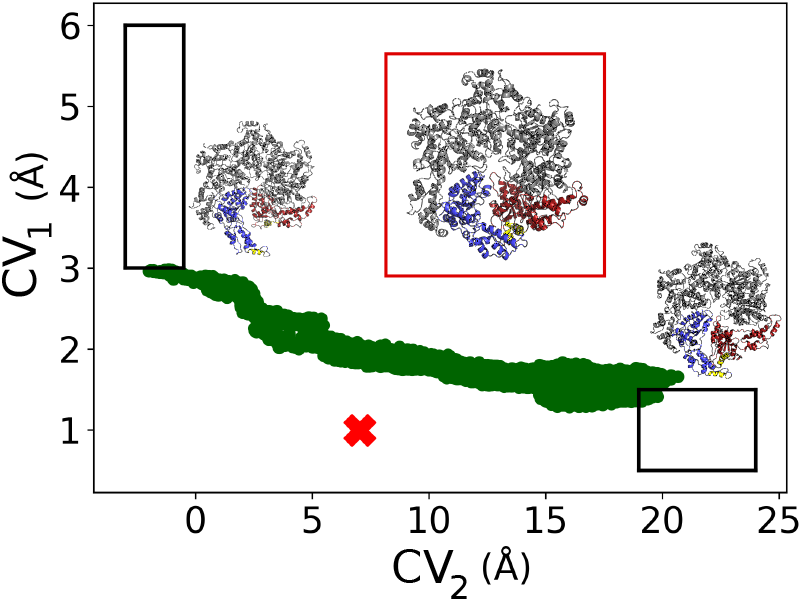
Katanin conformational space as a function of CV_1_ and CV_2_, as predicted by Ratchet&Pawl MD and WT-MetaD simulations (green region). The latter predicted also an intermediate outside that region (“X”). The boxes represent the spiral (upper left), intermediate (middle) and ring (lower right) structures, colored as in Fig. 2. The complete landscape from WT-MetaD is plotted in Fig. S2

**Fig. 4.**
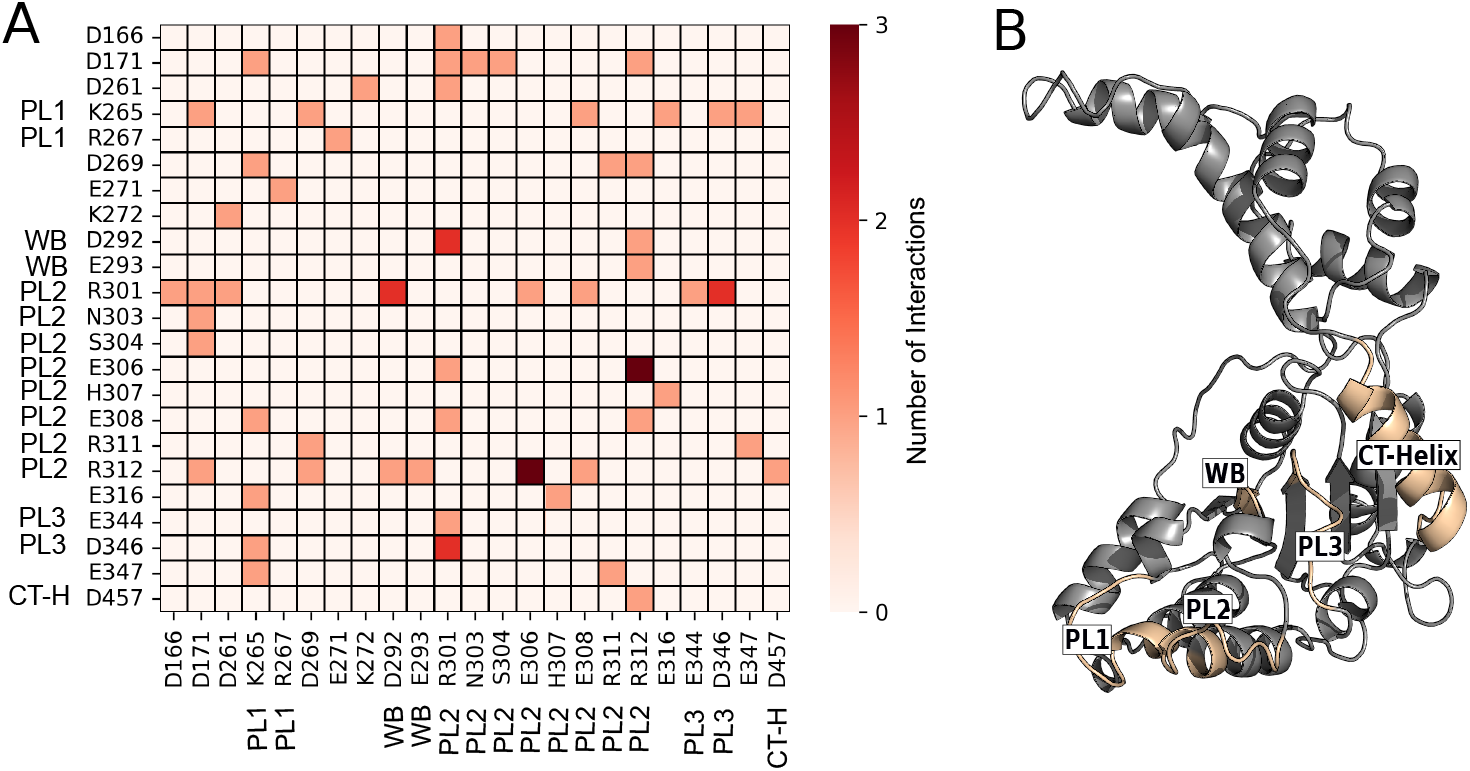
(A) Heatmap displaying the salt bridges/hydrogen bonds included in CV_2_). Residues that make at least one interaction are listed. The number of interactions are indicated by the color bar. Residues that are a part of a functional region are labeled with the latter. (B) Katanin monomer with most functional regions included in CV_2_ labeled.

Finally, we performed multiple walker WT-MetaD simulations on four independent replicas for 700 ns each (2.8 *µ*s in total) to investigate the conformational space as a function of the two CVs.

These simulations predicted a complex conformational space associated with the S → R transition (Fig. S2). This includes not only the region identified by Ratchet&Pawl MD, but also a minumum located outside that region (“X” in Fig. 3). This minimum was not sampled by the Ratchet&Pawl MD. To see if this minimum represents really an intermediate in the transition, we ran a 100 ns MD simulation starting from a representative conformation of the minimum. We observed that our intermediate structure remained within the region of the minimum for the entire dynamics, confirming the presence of a stable intermediate (Fig. 3). The latter is a hexamer with a 40° twist angle, differing from the spiral and ring’s 60° twist (Fig. S6). The backbone of the intermediate deviated 5.8 Å and 6.1 Å compared to the spiral and ring conformer, respectively. Many contacts between the NBDs of protomers A and B present in the spiral conformer and in the intermediate are lost in the ring one (Fig. 3, Tab. S2). This makes sense since the severing event is associated with an increase of flexibility to reach the ring state, which is successfully captured in our simulations.

## 2. Discussion

We have presented a computational study of the S → R transition in katanin, a protein structurally and functionally similar to spastin, with the goal of uncovering possible roles of HSP-linked mutations for the transition. The conformational landscape associated with the latter has been predicted by Ratchet&Pawl MD and metadynamics, as a function of two CVs. While the first (CV_1_) is constructed by us (it is simply the distance between selected helices of the protein), the second (CV_2_) is built automatically based on MD simulations on the two conformers. CV_2_ turns out to incorporates a variety of geometrical features involved in the S → R transition. Strikingly, almost all of the features of CV_2_ (31 out of 32) turn out to involve six residues - conserved on passing from katanin to spastin (see Tab. S2) and with similar chemical environments (Fig. S3A-B) - which undergo disease-linked mutations (9). All of them (in bold face hereafter) turn out to play a functional role: **R312** from PL2 (R460 in human spastin) interacts with residues from the CT-helix. This interaction is essential for hexamer stabilization (9, 15, 26). **R301** (and the correspondent arginine in spastin) from PL2 plays a key role in CTT binding, ATP binding, and hexamer oligomerization in both spastin and katanin (9). It forms salt bridges with **D346** from PL3 - important for oligomerization (11) - and with **E293**, a residue in Walker B that coordinates the Mg(II) ion (Fig. S4) (9). **D292** is located within the Walker B motif of the NBD. Its disease-linked mutation to a glycine of the corresponding aspartate in spastin (9) affects the magnesium binding site and hence ATP hydrolysis. Finally, **R311** in PL2 facilitates interprotomer communication (11). The correlation between the disease-related mutations in spastin and the corresponding residues in CV_2_ for katanin is fully consistent with the fact that the corresponding residues in spastin are crucial for the function of the enzyme.

WT-MetaD simulations of the S → R transition as a function of the two CVs point to the presence of a previously unrecognized intermediate (Fig. 3). As many as eleven residues, conserved between katanin and spastin (Tab. S2) undergo disease-lined mutations in the latter, form stabilizing interactions in the intermediate. The chemical environments for all the residues are similar across the two proteins (data not shown). **K239**, located in the Walker A motif, forms a salt bridge with D292 of Walker B (Fig. S5) and it is necessary for ATP-binding. The salt bridge made between **E207** and a neighboring protomer’s **R414** stabilizes the hexamer, since the latter belongs to the longest *α* helix on the protomer’s concave interface that makes direct contact with its neighbor. E207 also makes intra-protomer interactions with **R223**, which is necessary for the oligomerization of the hexamer. Here, **D295** in the intermediate interacts with R301 in a neighboring protomer. This points to its role in oligomerization. **R275** is also crucial for katanin’s oligomerization into a functional hexamer (11). It forms a salt bridge with E271 that impairs ATPase activity when mutated. **E279**, in the NBD, forms a salt bridge with R282. The corresponding residue in spastin disrupts the folding and stability of the ATPase domain when mutated to a proline (9). **R367**, located in the HBD, forms an intra-protomer salt bridge with D363. This is essential for the folding of this domain. **R356** in the CT-helix form a salt bridge with D467. Finally, **R351** in the Arginine finger is necessary for ATP hydrolysis. It forms an inter-protomer salt bridge with D457 to coordinate the Arginine finger with the CT-helix of the neighboring protomer. The D292, E293, R301, and R311 disease-related residues play also a role in the intermediate’s stability. All of these interactions are present in either the spiral and/or the ring conformers of the two proteins. Finally, we notice that D292, E293, R301, and R311, which have already been noted, play also a role in the intermediate’s stability (Fig. 5). **F352**, in the Arginine finger forms a salt bridge with D292, the Walker B residue important for Mg coordination, in the neighboring protomer. This interaction makes sense since the assigned role from spastin mutations shows this residue activates ATP hydrolysis in the neighboring protomer. Since the ATP-hydrolysis is required for the S → R transition, it makes sense that the intermediate is characterized by interactions with this Arginine finger residue.

**Fig. 5.**
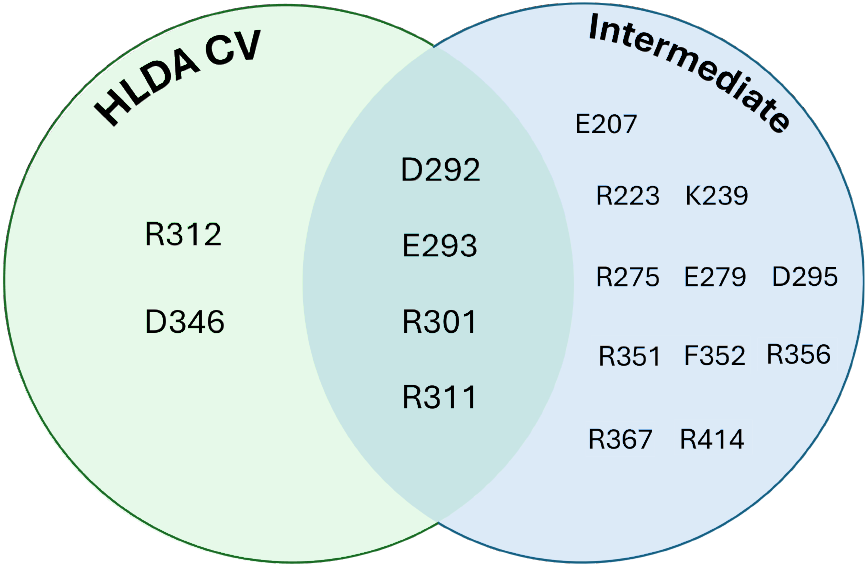
Summary of katanin residues part of CV_2_ and/or forming stabilizing interactions in the intermediate of the S → R transiton (Fig. 3), fully conserved in spastin and undergoing mutations associated with HSPs (see Tab. S1)

## 3. Conclusions

We have here defined a robust protocol that combines data-driven identification of relevant interactions with enhanced sampling methods to identify structural determinants of conformational transitions in large biomolecular complexes. Applying this protocol to katanin, (i) we ranked the most relevant non-bonded pairs from structural data using molecular simulation-based approaches and (ii) we identified an intermediate along the S → R transition which suggest a mechanistic interpretation for disease-related mutations which impair the function of severing enzymes (like spastin and katanin). Six disease-linked mutations affect the transition while eleven other positions that undergo mutations in disease are found to stabilize the intermediate (Fig. 5). This approach can be readily adapted to other large protein complexes, enabling the exploration of the system conformational space and the identification of transient states that are essential for function but challenging to capture experimentally.

### Materials and Methods

#### Katanin System Preparation

The two cryo-EM structures of the katanin complex, namely the spiral (6UGD, 3.5 Å) and ring (6UGE, 3.6Å) conformations, were obtained with ATP and a polyglutamate substrate in place of the microtubule tubulin tail (10, 11). Both structures contained missing residues in the ranges of 183-187 and 324-331 that were modeled using the Modeller program version 9.23 (27). To model the transition from spiral to ring, the nucleotide from protomer A was removed from the spiral to match the configuration of the ring. This aligns with katanin’s conformational transition, where stochastic cycles of ATP hydrolysis and removal from the hexamer result in increased flexibility within the terminal protomer that closes to the ring structure. In both conformations, we also modeled the TUBB sequence of *β*-tubulin (betaWT) in place of the solved polyglutamate substrate using PyMOL version 3.0 and GROMACS 2022 with the GROMOS 54a7 force field to construct and perform energy minimization of the betaWT sequence, respectively (28–32). This sequence was docked in place of the minimal substrate using the GRAMM protein docking sever from the Vakser lab by aligning the betaWT sequence to the minimal substrate in order to select the best starting position (33).

#### Molecular Dynamics Simulations

We ran molecular dynamics (MD) simulations of spiral and ring configurations described above using GROMACS 2022 (31, 32). The automated topology builder (ATB) was utilized for ATP parameters (34). Each structure (∼19,000 atoms) was placed in a rhombic dodecahedron water box to increase computational efficiency compared to the size of the cubic box, and the water molecules were represented with SPC-16 explicit solvent with 67 sodium ions to neutralize the total charge of the system (35). Energy minimization was done with the steepest descent algorithm and the Verlet cutoff-scheme (36) for 50,000 steps. For NVT and NPT equilibration, we used the velocity-rescaling thermostat at 300K with the leap-frog integrator algorithm and the Parrinello-Rahman barostat at 1.0 bar, respectively (37, 38). Each equilibration step was run for 500 ps, with a thermostat coupling frequency of 0.1 ps and a barostat coupling frequency of 2.0 ps. The short-range nonbonded interactions were calculated using a distance cutoff of 10.0 Å and a dispersion correction for energy and pressure for anything past the cutoff. Particle Mesh Ewald was used for long range electrostatic interactions (39). The LINCS method was used for the bond lengths involving hydrogen atoms, and the simulations were run using a 2 fs integration step (40). For each system, we ran for a total of 150 ns and checked the convergence of the trajectories using the backbone RMSD, where the first frame of the production run was used as the reference frame (41).

#### Harmonic Linear Discriminant Analysis

To identify a minimal number of apt collective variables (CVs) which can efficiently represent the spiral-ring transition, we employed a protocol previously successful in capturing complex biological changes within fewer dimensions using harmonic linear discriminant analysis (HLDA) (21, 22).

This approach starts with a set of observables (called local descriptors) whose distributions in the states of interest are studied. From the analysis of these distributions, a linear combination of all these local descriptors is generated that maximizes the separation of the states. For HLDA, the within-class scatter matrix (*S*_*w*_) used to measure the degree of class separation is calculated using the harmonic average to place more weight on states with smaller variances, where (Σ_*A*_, Σ_*B*_) is the multivariate variance for each metastable state (21).

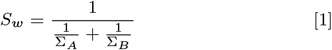

The collective variables using HLDA (*s*_HLDA_(*R*)) is then calculated in eq. 2, with *R* representing atomic coordinates, *µ*_*A,B*_ the expectation values per state, and *d*(*R*) the local descriptors.

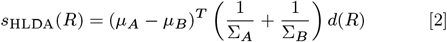

Here, to have an insight on the interactions that regulate the stabilization of the two states, we collected unique salt bridges and hydrogen bonds between the spiral and ring using unbiased simulations in the ring and spiral states. VMD was used to collect the salt bridges and hydrogen bonds (42). Salt bridges were calculated using an oxygen-nitrogen cut-off of 6 Å in attempts to collect more critical differences between the conformations (25). For the hydrogen bonds, we set a threshold for the donor-acceptor distance of 3.0 Å with a cutoff angle of 30 degrees. We only considered inter-protomer salt bridges and hydrogen bonds except for the two terminal protomers, where we additionally included intra-protomer interactions.

#### Ratchet&Pawl Simulations

To test the optimization of our selected CVs and to compare with later metadynamics simulations, we used Ratchet&Pawl MD (rMD) to model katanin’s transition from spiral to ring (18, 19). rMD is a non-equilibrium technique: after the definition of an apt CV and two states, we can define a transition direction (i.e., we define the starting and the ending states) and we apply an harmonic potential which disfavor the system in going in the opposite direction with respect to the desired one. The harmonic potential term on the CVs helps the system transition between two states. Such harmonic potential follows the system during the transition, staying still when the system tries to come back to the initia tate along the CV. The potential is thus

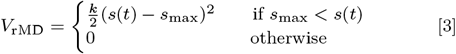

Where *s*_max_ is the maximum value of the CV during the simulation and *k* is the force constant. Considering this, rMD allows us to study katanin’s transition path between a starting and end state. To describe the hinge-like motion of the terminal protomers in the severing event, we calculated the center of mass distance between an alpha helix in protomer A (residues 199-215) and protomer F (residues 425-436) (3, 9). These helices were selected based on giving the most extreme distinctions between the spiral and ring state as well as their direct path to forming interactions with one another in the ring. We added another CV using the 1st eigenvector of the HLDA analysis, using only the top 32 salt bridges or hydrogen bonds that had the largest contribution in distinguishing spiral and ring while also interacting with one of the three functional pore loops to reduce the “noisy” elements that would decrease simulation efficiency while also retaining the connection between katanin’s involvement with the microtubule substrate through the pore loops with the severing motion seen through the spiral to ring transition. We ran 10 simulations of the forward (spiral to ring) and backward (ring to spiral) transitions until katanin had reached the CV boundaries defined in the initial MD runs, which took around 60-70ns to obtain each transition. All CV calculations and rMD were done via GROMACS 2021.6 patched with PLUMED 2.8 (43, 44). The harmonic constant for both CVs was fixed to k = 500 kJ/mol/nm^2^.

#### Metadynamics Simulations

We ran WT-Metad simulations using the two CVs from our rMD runs (23, 24). We performed an initial simulation to survey katanin’s transition from spiral to ring. After 400 ns, we selected 4 configurations from such run to use for a Multiple-Walkers Well-Tempered Metadynamics run (45). These include the spiral and obtained ring structures along with two structures found on opposing sides of the obtained landscape, to maximize the exploration of conformational space. The bias factor set was 20, and the height of the Gaussians were 1.2 kJ/mol, deposited at a pace of 500 steps. The widths of each Gaussian deposited were 0.05 for each of the two CVs. The temperature was set to 300 K. We included upper and lower repulsion potentials using the unbiased MD runs of the spiral and ring states as boundaries for each CV since the calculated HLDA CV is only relevant in describing the spiral-to-ring transition. The multiple walker simulations continued until the combined total simulation time reached 2.8 *µ*s (700 ns per walker).

#### New intermediate molecular dynamics simulations

To test the stability of the intermediate state identifield from the multiple-walker WT-MetaD simulations, we tested it with plain MD simulations. We selected the starting structure for this run by taking the centroid from the k-means cluster of the WT-MetaD space that contained only the minimum region. We thus ran 100 ns of plain MD, verifying that the system did not exit the minimum, with the same simulation condition detailed in the previous sections.

## A. Data Availability Statement

All the data and input files needed to reproduce these simulations, as well as trajectory files are available on the public repository Zenodo, with the identification code 10.5281/zenodo.14218204.

## ACKNOWLEDGMENTS

This project used computing time granted through NHR4CES on CLAIX-2018. Additional computational support came from the advanced computing systems available through ACCESS based on the allocation BIO210094 (to RID). This research was funded by the National Science Foundation MCB-1817948 (to RID).

## Notes

https://zenodo.org/records/14218205

### Competing Interest Statement

The authors have declared no competing interest.

