## Supplemental Information for "Structural and functional role of residues undergoing hereditary spastic paraplegias-linked mutations: insights from a simulation of the spiral to ring transition in katanin"

### Introduction

#### 1. Structural Differences Between Katanin and Spastin

Katanin and spastin, despite being strongly related, differ in the following structural determinants: (i) katanin has a 'fishhook' motif required for interface interactions between the protomers and spastin does not. (ii) katanin has two subunits (catalytic p60 and regulatory p80), while human spastin is a homomeric enzyme. (iii) human spastin has a dedicated microtubule-binding domain (MTBD), while katanin relies on the p60 subunit for severing and the p80 subunit for regulation. Katanin's p80 subunit and spastin's MTBD are more important concerning directing and enhancing MT binding.

The ATPase motors between the severing enzymes contain the same functional elements and are composed of an NBD and HBD (*see main text*). However, the cryo-EM structures of their spiral states (katanin, *C.elegans* and spastin, *D. melanogaster*) show differences in sequence length and secondary structure. There is a 1.13 Å deviation between the backbone of each protein's protomer A. Katanin has 316 residues per monomer while spastin has 303 residues (**Chart S1)**. Both domains are slightly smaller in spastin, with 25 and 24 secondary structures in the NBD and 11 and 9 secondary structures in the HBD for katanin and spastin, respectively. A larger structural difference is at the HBD tip (denoted by * in the figure below), where spastin's HBD tip shows a short helix while katanin has a longer flexible loop going into a helix. A previous studies from some of us found this region in katanin to be significantly more flexible than in spastin, which could influence the stability in the hexamer due to the HBD's role in forming protomer-protomer interfaces.[1]

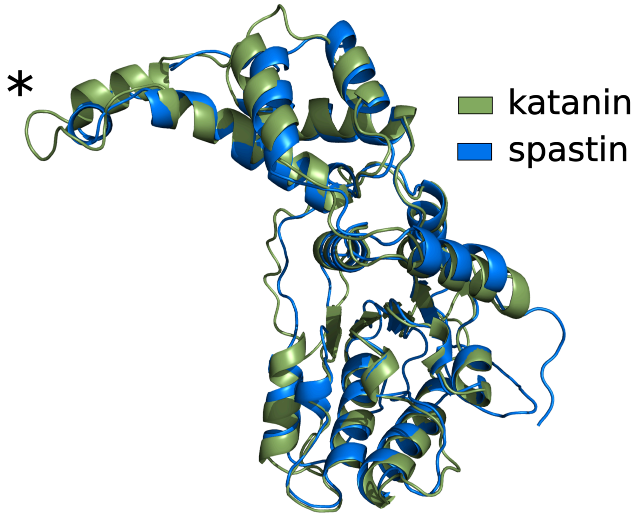


**Chart S1**

### 2. Katanin and Spastin: a comparison among structural determinants and function.

We carry out the calculations on katanin from *C. elegans* instead of human spastin, for several reasons: **(i)** both the spiral and the ring conformations are available only for katanin[3]. **(ii)** The differences between the two proteins are expected not to impact the spiral-ring transition because the different region's functional role is to direct and enhance MT binding.[4,5] **(iii)** The fold, the functional regions, and the vast majority (78%) of the residues undergoing disease-linked mutations are fully conserved across the two proteins (Tab. S2), [6] with very similar chemical environments (as shown by visual inspection) (data not shown).. **(iv)** katanin can partially compensate for the HSP-related loss of spastin activity in neurons, suggesting a very similar functional role *in vivo* of the two proteins.[7]

### 3. Figures

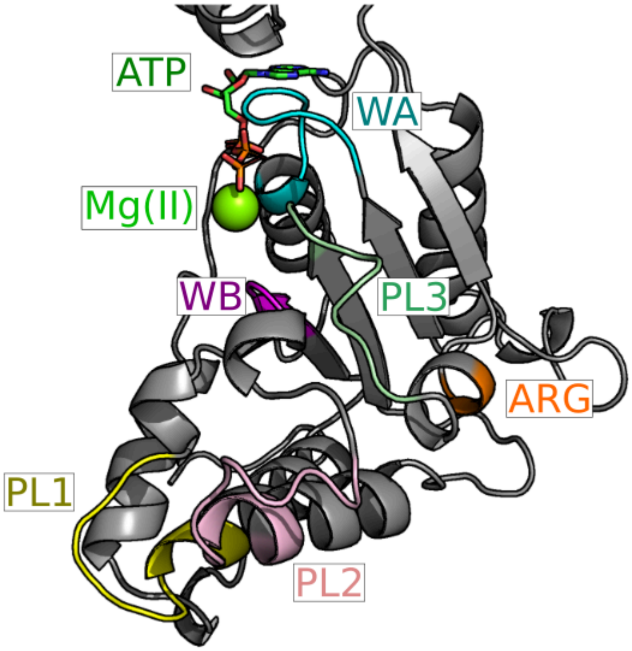


Fig S1. View of katanin NBD detailing the ATP and CTT binding sites.

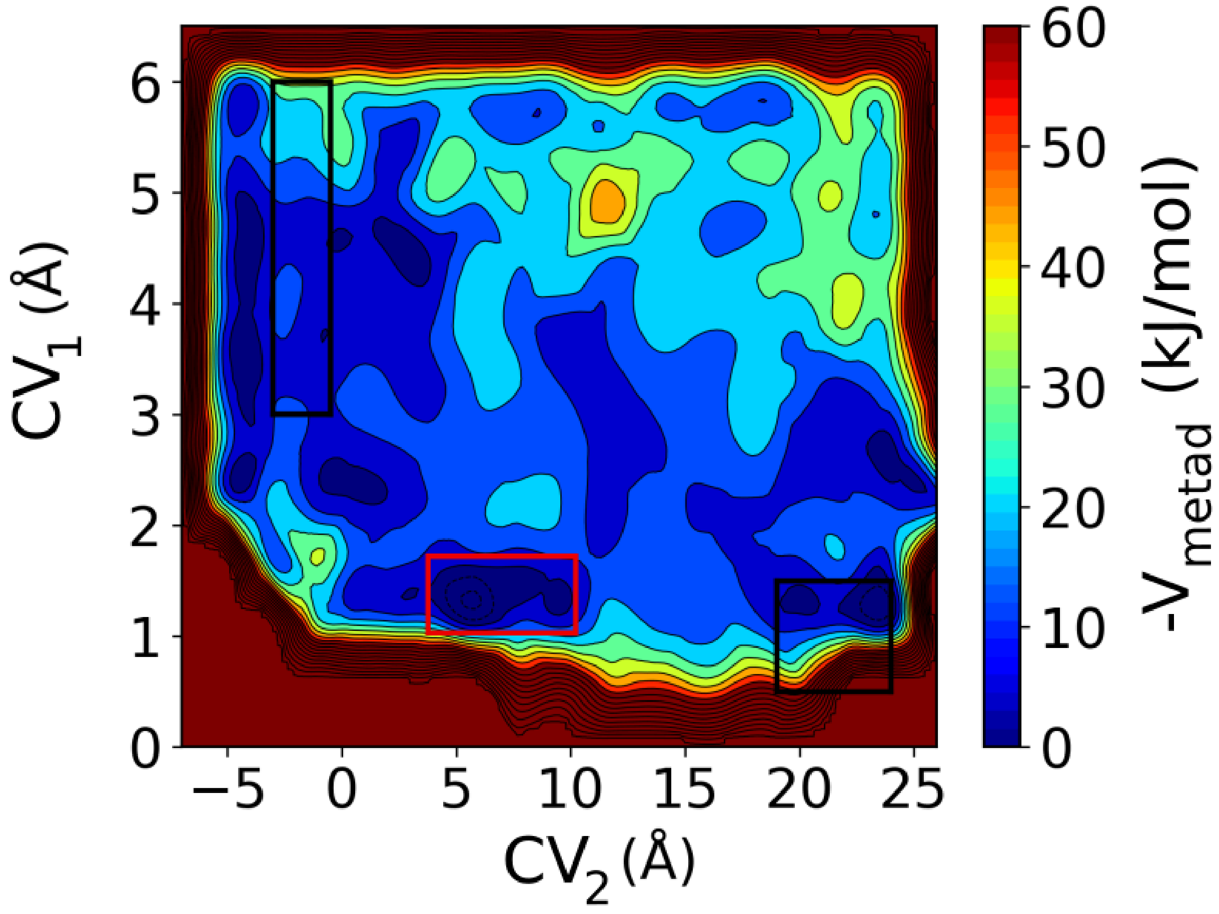


Fig S2. Negative potential deposited during multiple-walker well-tempered Metadynamics simulation. This plot provides a qualitative estimation of the part of the conformational space (projected along the CVs) which are accessible during simulations. The regions in dark blue color between the two stable states (black rectangles) are conformations which are likely to be visited during the S → R transition. The spiral and ring state boxes are defined similarly to Fig. 3 in the main text. The intermediate region is indicated by the red box.

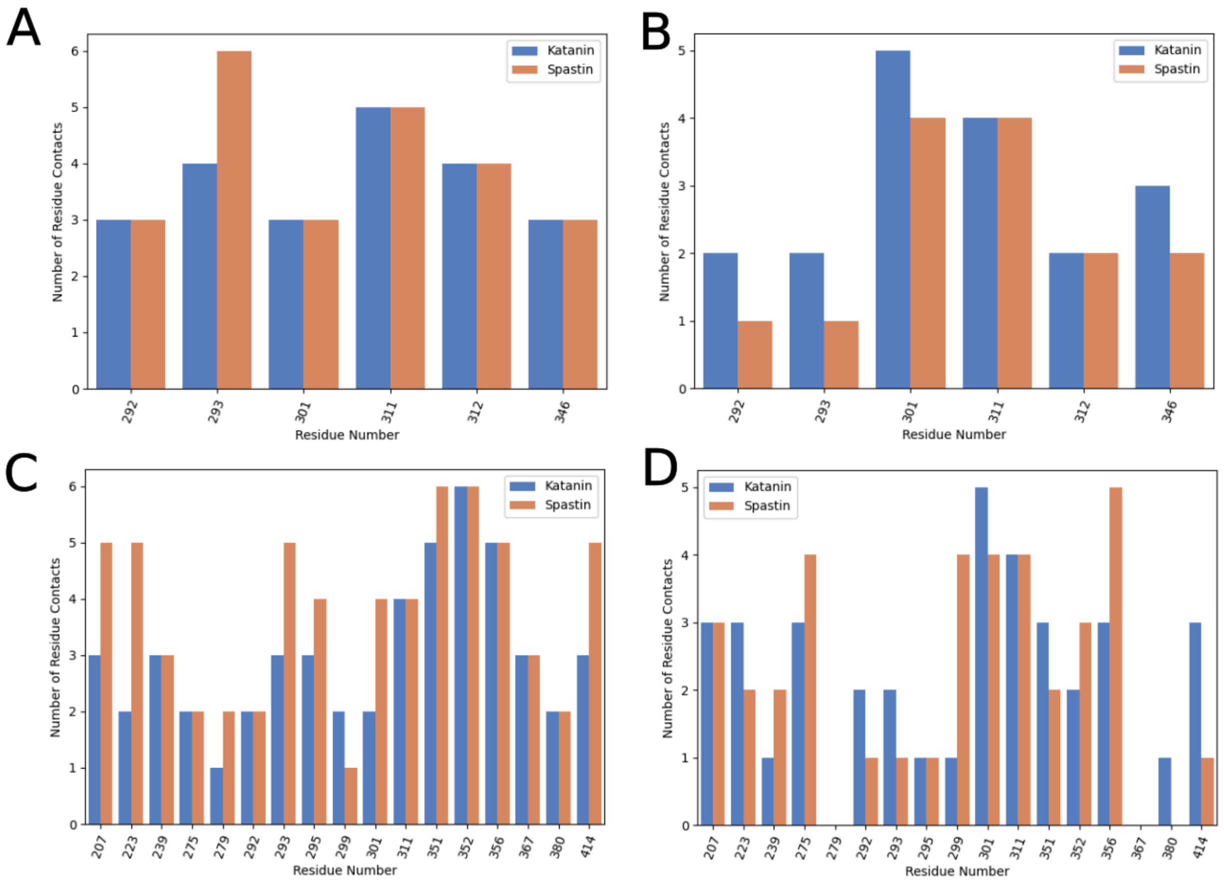


Fig S3. The number of (A,C) **intra** and (B,D) **inter**-protomer contacts calculated for disease-related residues in spastin and aligned residues in katanin. Top row shows residues found in $CV_2$ as important for the S→R, and bottom row shows residues important for the stability of the intermediate. Protomer C of each protein's starting configuration was analyzed to compare the environment of the hexamer where a protomer has neighbors along both interfaces. Katanin residues are used to label the x-axis. Spastin alignments are listed in Table S1. Contacts for a residue reports the number of unique residues for which at least one pair of heavy atoms, one from the central residue and one from the partner residue, were within 8 Å. The calculations were carried out using the MDAnalysis Python package.[2.]

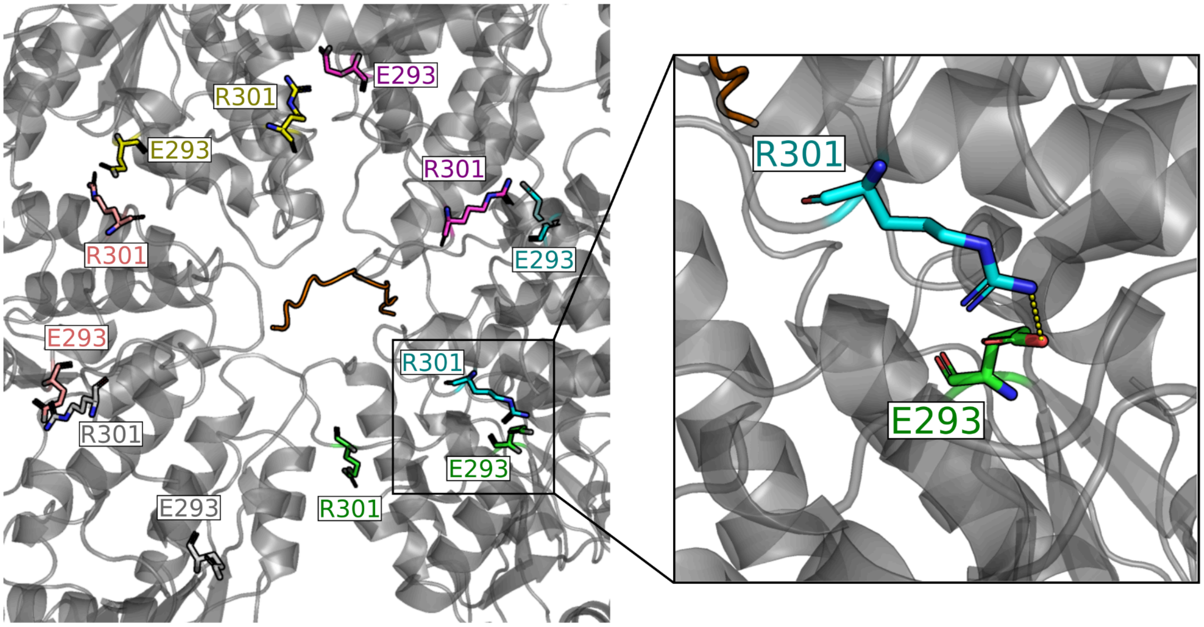


Fig S4. Interactions between R301 and E293 that are in salt bridge distance between protomers.

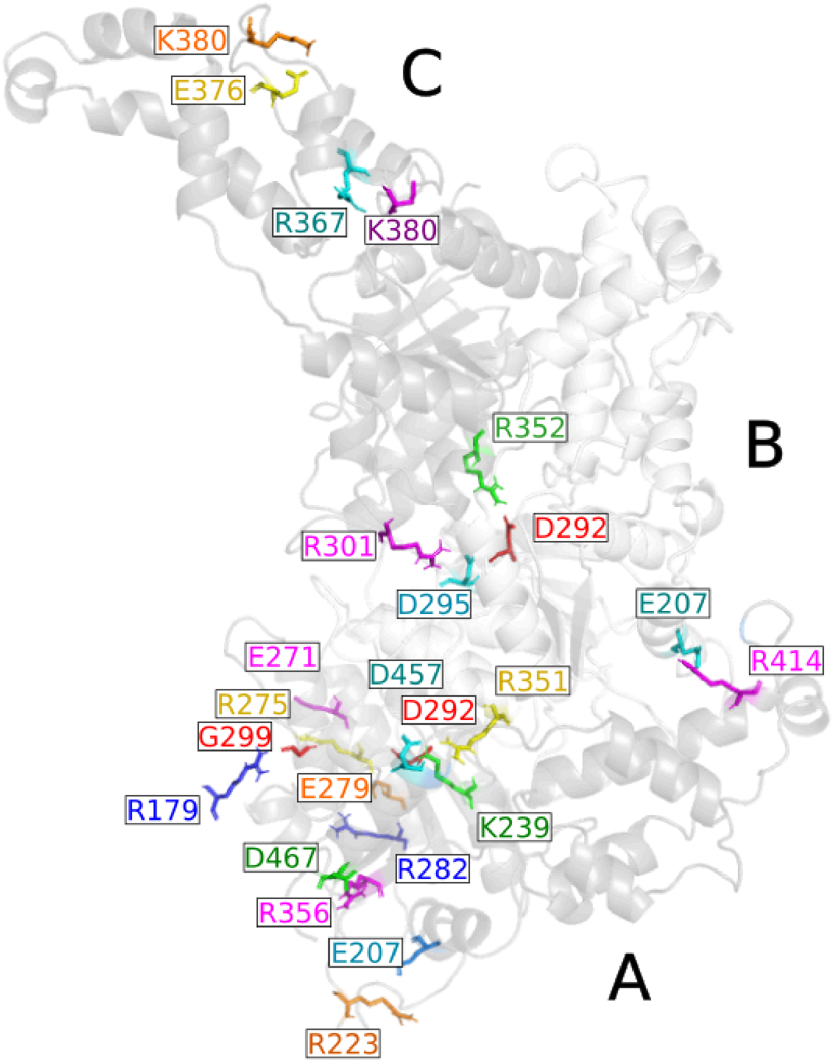


Fig S5. Salt bridges/hydrogen bonds found specifically important for the stability of katanin's intermediate. Protomers A, B, and C shown.

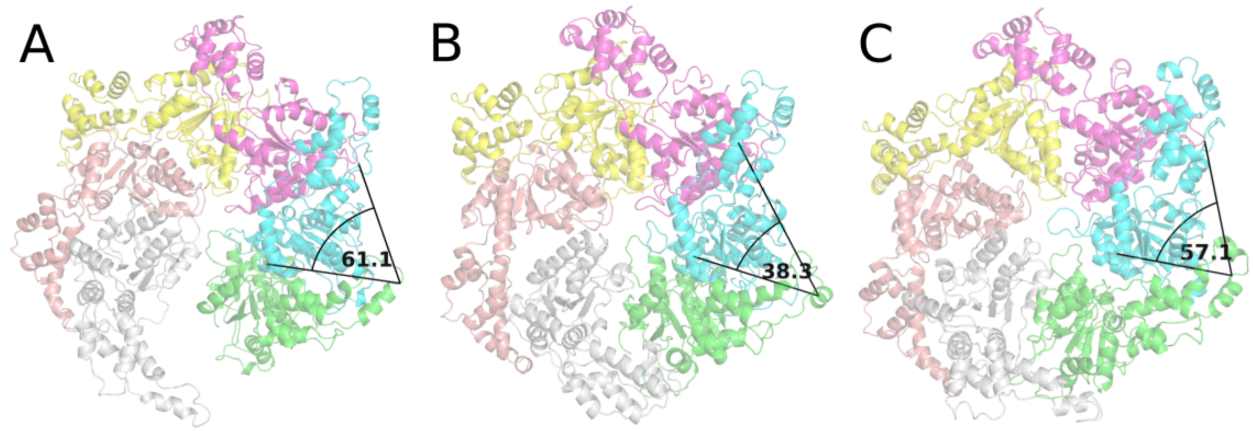


Fig S6. Protomer twist angle of (A) spiral (B) intermediate and (C) ring state.

### Tables

| Spastin Residue undergoing mutation(s) | HSP-linked mutations | Impaired Function when Mutated | Corresponding residue in Katanin |
|---|---|---|---|
| **I344** | K | Nucleotide Binding | **I195** |
| **G346** | C | Nucleotide Binding | **G197** |
| Q347 | K, P | Nucleotide Binding | M198 |
| **E356** | G | Nucleotide Binding | **E207**[#] |
| **L360** | P | Oligomerization | **L211** |
| **P361** | L, S | Oligomerization | **P212** |
| S362 | C | - | R356 |
| R364 | M | Oligomerization | V215 |
| L367 | S | Oligomerization | F218 |
| **F368** | L, S | Oligomerization | **F219** |

| Spastin Residue undergoing mutation(s) | HSP-linked mutations | Impaired Function when Mutated | Corresponding residue in Katanin |
|---|---|---|---|
| T369 | P | Oligomerization | Q220 |
| **G370** | R | Oligomerization | **G221** |
| **R372** | G | Oligomerization | **R223#** |
| **P374** | R | Oligomerization | **P225** |
| G377 | E | Folding & Stability | A228 |
| L378 | G | Folding & Stability | M229 |
| F381 | C, L | Folding & Stability | A232 |
| **G385** | E | Nucleotide Binding | **G236** |
| N386 | K | Nucleotide Binding | T237 |
| **K388** | R | Nucleotide Binding | **K239#** |
| M390 | V | Nucleotide Binding | L241 |
| **S399** | L | Folding & Stability | **S250** |
| I406 | V | Folding & Stability | V257 |
| **S407** | R | Oligomerization | **S258** |
| **G417** | E | Oligomerization | **G268** |
| **V423** | L | Folding & Stability | **V274** |
| **R424** | G | Oligomerization | **R275#** |
| **L426** | V | Folding & Stability | **L277** |
| **F427** | C | Folding & Stability | **F278** |
| A428 | P | Folding & Stability | E279# |
| **P435** | T, L | Folding & Stability | **P286** |
| **S436** | F | Folding & Stability | **S287** |
| **D441** | G | Mg Coordination | **D292*#** |
| **E442** | K | Nucleotide Binding | **E293*#** |

| Spastin Residue undergoing mutation(s) | HSP-linked mutations | Impaired Function when Mutated | Corresponding residue in Katanin |
|---|---|---|---|
| **D444** | E, N, G | Oligomerization | **D295**[#] |
| C448 | Y | Oligomerization | G299[#] |
| **R450** | S | Oligomerization | **R301**[*#] |
| **S458** | R | Folding & Stability | **S310** |
| **R459** | T | Oligomerization | **R311**[*#] |
| **R460** | C, L | Oligomerization | **R312**[*] |
| L461 | P | Folding & Stability | V313 |
| **L466** | V | Folding & Stability | **L318** |
| **G471** | D | Folding & Stability | **G323** |
| **N487** | D | Oligomerization | **N340** |
| **P489** | L | Folding & Stability | **P342** |
| **D493** | G | Folding & Stability | **D346**[*] |
| **R498** | S | Phosphate Binding | **R351**[#] |
| **F499** | C, H | ATP Hydrolysis | **F352**[#] |
| **R503** | L, G | Oligomerization | **R356**[#] |
| **R514** | G | Folding & Stability | **R367**[#] |
| G527 | N | Folding & Stability | K380[#] |
| **L537** | P | Folding & Stability | **L389** |
| **G546** | R | Folding & Stability | **G398** |
| L549 | P | Folding & Stability | V401 |
| A551 | Y, P | Oligomerization | S403 |
| D555 | N, G | Oligomerization | T407 |
| **A556** | V | Folding & Stability | **A408** |

| Spastin Residue undergoing mutation(s) | HSP-linked mutations | Impaired Function when Mutated | Corresponding residue in Katanin |
|---|---|---|---|
| **A557** | V | Folding & Stability | **A409** |
| G559 | D | Oligomerization | N411 |
| R561 | G | Oligomerization | L413 |
| **R562** | Q | Oligomerization | **R414#** |
| N579 | H | - | L437 |
| D584 | H, V | - | I441 |
| **W607** | C | Folding & Stability | **W465** |
| D613 | H | Oligomerization | A471 |

Tab S1. Residues of human spastin whose mutations (one to three) are associated with HSP. Aligned residues for katanin from C. elegans are listed next to them. The role of each residue determined in experimental mutation studies is listed if known. * indicates that the residue is included in $CV_2$ (Tab. S2). # indicates that the residue was found important in the Intermediate along the S→R transition. The residues in boldface are conserved across spastin and katanin.[6,8]

| Number of interaction | Interaction | Protomers |
|---|---|---|
| 1# | H307 - E316 | A - B |
| 2# | R301 - D261 | A - F |
| 3# | E306 - R301 | A - B |
| 4# | E271 - R267 | A - B |
| 5# | D346 - K265 | A - F |
| 6# | D166 - R301 | F - A |
| 7# | D171 - R301 | F - A |
| 8# | D171 - R312 | F - A |

| Number of interaction | Interaction | Protomers |
|---|---|---|
| 9[#] | D261 - K272 | A - B |
| 10[#] | E347 - K265 | B - A |
| 11[#] | D346 - R301 | A - B |
| 12[#] | E306 - R312 | A - B |
| 13[#] | D346 - R301 | F - A |
| 14[#] | E308 - R312 | E - D |
| 15[#] | D292 - R301 | A - B |
| 16[#] | E306 - R312 | B - C |
| 17[#] | E316 - K265 | B - A |
| 18[#] | D269 - R312 | A - A |
| 19[#] | D457 - R312 | C - D |
| 20[#] | D292 - R312 | C - D |
| 21[#] | E308 - K265 | F - F |
| 22[#] | D171 - K265 | B - A |
| 23[#] | D292 - R301 | E - F |
| 24[#] | E293 - R312 | A - B |
| 25* | S304 - D171 | F - A |
| 26[#] | D269 - D265 | B - A |
| 27[#] | E308 - R301 | C - C |
| 28* | D171 - N303 | F - A |
| 29[#] | E347 - R311 | F - F |
| 30[#] | E306 - R312 | D - E |
| 31[#] | E344 - R301 | B - B |
| 32[#] | D269 - F311 | F - A |

Tab S2. Intra- or Inter-protomer residue-residue H-bond (*) and salt bridge (#) interactions included in $CV_2$. The protomers A-F are shown in Fig. 2.
